# Dimensionality reduction and statistical modeling of scGET-seq data

**DOI:** 10.1101/2022.06.29.498092

**Authors:** Stefano de Pretis, Davide Cittaro

## Abstract

Single cell multiomics approaches are innovative techniques with the ability to profile orthogonal features in the same single cell, giving the opportunity to dig more deeply into the stochastic nature of individual cells. We recently developed scGET-seq, a technique that exploits a Hybrid Transposase (tnH) along with the canonical enzyme (tn5), which is able to profile altogether closed and open chromatin in a single experiment. This technique adds an important feature to the classic scATAC-seq assays. In fact, the lack of a closed chromatin signal in scATAC: (i) restricts sampling of DNA sequence to a very small portion of the chromosomal landscapes, substantially reducing the ability to investigate copy number alteration and sequence variations, and (ii) hampers the opportunity to identify regions of closed chromatin, that cannot be distinguished between non-sampled open regions and truly closed. scGET-seq overcomes these issues in the context of single cells. In this work, we describe the latest advances in the statistical analysis and modeling of scGET-seq data, touching several aspects of the computational framework: from dimensionality reduction, to statistical modeling, and trajectory analysis.

## 1 Introduction

Single cell Genome and Epigenome by Transposase sequencing (scGET-seq) has been recently developed to study the genome and epigenome from the same single cells [1]. scGET-seq introduces a biased tn5 transposase hybrid (tnH) which is able to target condensed heterochromatin exploiting the affinity of HP1-*α* chromodomain for histone modification H3K9me3. In the original publication we showed how scGET-seq could be used to get insight about the genetics of heterogeneous cancer samples as well as to derive cell trajectories by the analysis of changes in chromatin status. However, the methods used to process data were overly complicated. In fact, we obtained low-dimensional representations by using matrix tri-factorization [2] of four count matrices. At time of writing, the data fusion method we adopted is no more developed or maintained, which prompts to the pursuit of an alternative method. In addition, we used to count tn5 and tnH reads over two sets of genomic regions, representative of accessible and compact chromatin, based on the DnASE I Hypersentitive Sites census available for human [3] or mouse [4]. Since the number of regions in both sets are different, we needed to obtain a low-dimensional representation (20-dimensional UMAP) of each count matrix before data fusion, at the risk of introducing distortion artifacts.

To model cell trajectories we hacked the `scvelo` package [5], exploiting its routines developed to model RNA velocity [6]. To do so, we arbitrarily fed the algorithm with two matrices, derived from tn5 and tnH counts, in place of expected spliced and unspliced mRNA counts. In the original publication, we applied this approach to study human Induced Pluripotent Stem Cells (IPS) differentiating to Neural Precursor Cells (NPC). While we obtained biologically sound results (*i.e.* the selected regions were indeed associated to signatures of nerual differentiation), the approach has several critical points. RNA velocity is defined for a two-step process going from RNA synthesis to RNA degradation, a path that does make little sense from chromatin folding/unfolding, where changes are in principle reversible. Consequently, any model of RNA-velocity assumes a direction, a premise that does not hold for chromatin, although specific models might be generated. For example, it has been shown that committed cells have a globally more condensed chromatin conformation [7].

To address these points, we introduce two improvements of scGET-seq analysis workflow, enabling a more straightforward approach to data processing. First, we present a simple and effective strategy to count and integrate data from both the transposases. Second, we propose a strategy to derive cell trajectories and analyze dynamic changes in chromatin accessibility abandoning the inappropriate RNA-velocity approach.

## 2 Results

### 2.1 Embedding scGET-seq data

To simplify the analysis procedure, we generate only two count matrices instead of four, one for each transposase. Each count matrix only reports the number of aligned reads, without duplicates, that fall into fixed-size genomic bins. We chose 5 kb as default size, as previously done in [8]. This step generates two equally sized matrices. After filtering out cells with low coverage and regions with excessive zero-count in both matrices, we normalize data according to the total coverage and apply log-transformation. The next step is to derive a single low-dimensional embedding representing both enzymes at the same time. While many approaches have been developed for analysis of multiomic data [9], we observed that our data could be intuitively represented as a tensor, so that we could apply a tensor decomposition strategy. Among the available approaches, we chose Tensor-Train Decomposition (TTD) [10], mainly because of its scalability and speed. By using TTD, data processing becomes conceptually similar to Singular Value Decomposition (SVD), largely used to reduce dimensionality in scATAC-seq data [11].

We show the effectiveness of such approach analysing the Caki/HeLa cell mixture reported in [1]. There, a mixture of two cell lines in known proportions (80% Caki-1, 20% HeLa) have been profiled by scGET-seq. We retrieved the publicly available data (E-MTAB-9650) and performed the entire analysis from the beginning. We compare TTD embedding with PCA embedding, calculated on tn5 counts, and the original data fusion approach (hereby called “Fusion”). We derived a *k* NN graph using the same number of neighbors and identified cell groups by stochastic block models using `schist` [12], so that cell groups are consistently called in three datasets. We then evaluated the ability of such graph to describe cell identities, as indicated in the original dataset, in terms of cluster completeness (*i.e.* cell types correspond to inferred cell groups), homogeneity (*i.e.* inferred cell groups only contain cells from a single cell type) and V-measure (harmonic mean of both scores). In Figure 1 we show the UMAP produced from three different embeddings, colored by cell type or the highest level in group hierarchy. Cell groups obtained after PCA embedding tend to produce overclustered data, although HeLa cells are still identifiable (cluster 3). Fusion and TTD embeddings, instead, allow cell groupings consistent with the ground truth.

**Figure 1:**
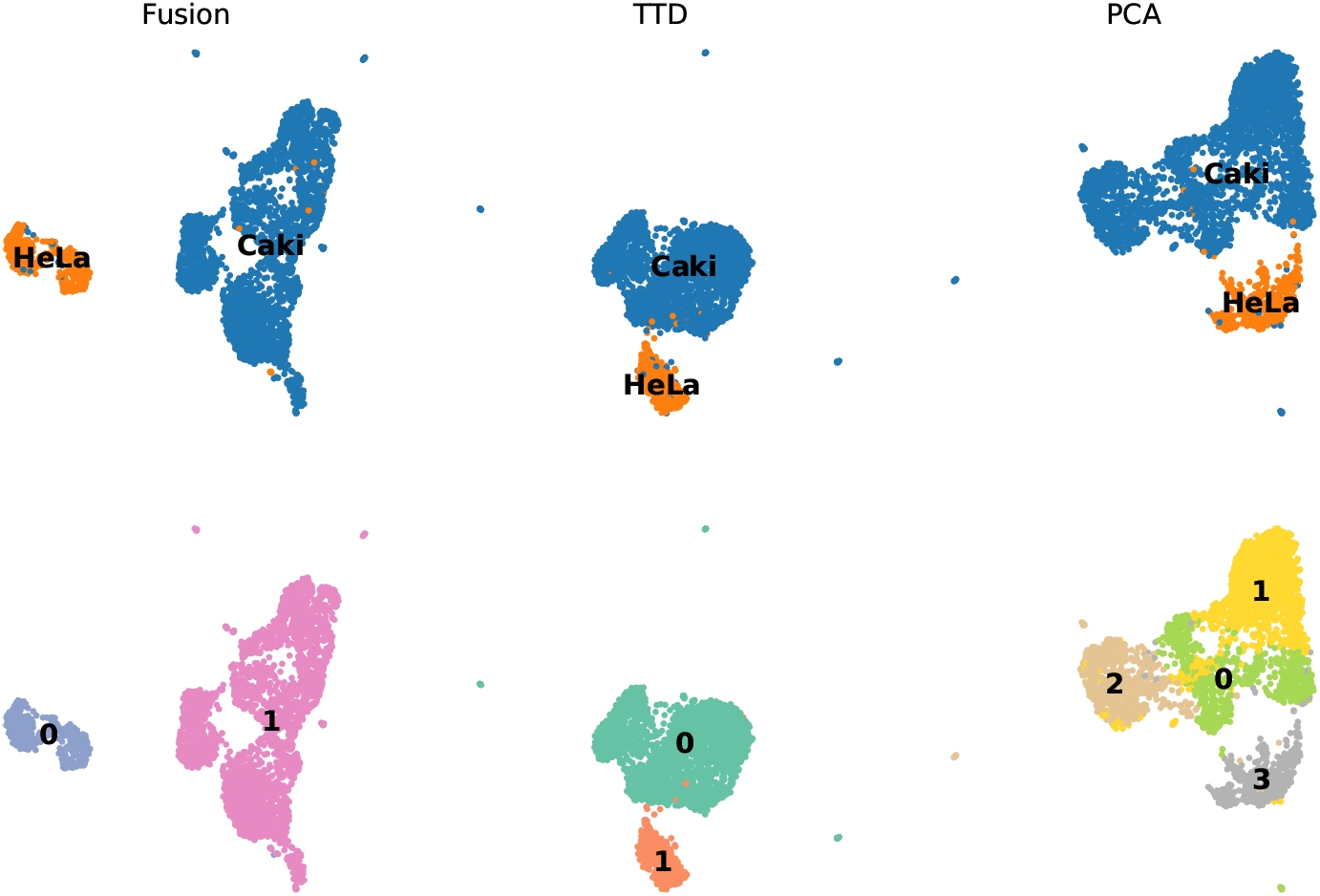
UMAP embeddings representing a mixture of Caki/HeLa cells obtained with three different dimensionality reduction methods. Each column correspond to a different method (from left to right: Matrix Tri-factorization, Tensor Train Decomposition, Principal Component Analysis). In the top row cells are colored according to cell identities, in the bottom row cells are colored according to the higher level of the nested stochastic block model provided by `schist`.

Consequently, the definition of cell groups performs worst when PCA is used (Table 1). Given that TTD is conceptually simpler, it is faster and, most important, the package used to perform matrix tri-factorizaion [2] is not actively maintained, we argue that TTD is an effective strategy to produce embeddings of scGET-seq data.

**Table 1:**
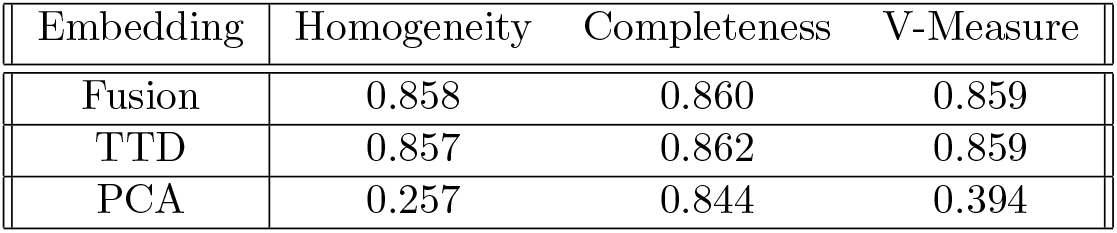
Performance of different embedding methods in consistently identify cell groups that correspond to ground truth. For each method we report completeness, homogeneity and V-measure

### 2.2 Scoring accessibility

TnH and tn5 have indeed different enrichments in accessible or compact chromatin and the difference between two signals could indicate chromatin status [1]. In principle, the plain difference between two signals could be explicative enough of the underlying chromatin status. However, a proper measure should take into account the total and relative coverage of both enzymes, that can vary across cells. Also, while tnH has a strong tropism to heterochromatin, it also retains a marginal activity in accessible regions. At the same time, aspecific signal from tn5 is distributed over the entire genome, to the extent it can be used to call copy number alterations at coarse resolutions [13]. Therefore, a proper measure of net accessibility should also consider cross-domain contamination. A convenient way to address this problem is to treat read counts in each bin as random variables and then associate the probability to observe a specific difference as proxy of accessibility.

scATAC-seq data are usually modelled according to Poisson law [8], a distribution that is also suitable for signals spread along the genome [14], as in the case of tnH. We hence posit that both tn5 and tnH can be modelled using Poisson law, possibly introducing Zero Inflation correction to address the high ratio of drop-outs. Observed counts in each bin are treated as random variables, *X*_5_ and *X_H_* for tn5 and tnH respectively, and accessibility is defined as the probability associated to the tail of the cumulative density function of *X*_5_ *− X_H_*. Given

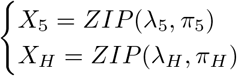

and *t* being the smallest integer so that

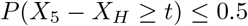

then accessibility *A* is defined as

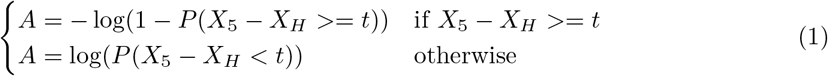

so that it is proportional to the excess of one transposase. Typically, the probability of observing 0 counts in one or the other transposase is high, therefore *P*(*X*_5_ *−X_H_* = 0) is also high, due to unobserved counts. For the same reason, this formulation results in a bias to positive values of *A*. We hence decided to set *A* = 0 when both tn5 and tnH are empty. If counts were modelled using a Poisson law, the difference would follow a Skellam distribution. In case of ZIP distribution, there’s no closed form to define the distribution of the difference, therefore we calculate *p*-values using the empirical cumulative density function.

To exemplify our implementation, we took advantage of a published experiment (E-MTAB-9651) in which NIH-3T3 cells have been treated with a short-hairpin RNA to reduce the expression of Kdm5c (sh); in the same experiment, a batch of cells is treated with scramble RNAi, as control (scr). As previously reported, reduction of Kdm5c expression causes major rewiring of pericentromeric heterochromatin [15], in addition to the increase in genomic instability. Another interesting property of such experiment is that downregulation of Kdm5c causes impaired cell cycle with prolonged G1 phase and a delay of entry in S phase [16].

First of all, we tested if there is sufficient evidence of using ZIP in place of Poisson using a Likelihood Ratio Test. For each cell we calculated the mean of per-bin coverage, excluding mitochondrial reads, as Maximum Likelihood Estimator of *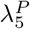* and 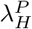, the only parameter defining Poisson distribution for tn5 and tnH respectively. At the same time, we infer by MLE 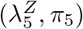 and 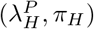, the expected coverage and the expected fraction of zeros for each cell. In all, our data suggest that ZIP is appropriate to describe both tn5 and tnH, having overall the highest FDR corrected *p*-value equal to 1.016 × 10^*−5*^ for tn5. For practical purposes, however, we decided to fit ZIP using the Method of Moments which is slightly faster [17].

After evaluating *A* for each region in each cell, we derive a global rate of chromatin compaction or relaxation calculated as

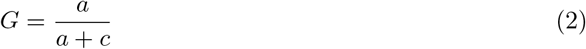

where *a* is the number of regions having *A >* 0 and *c* is the number of regions having *A <* 0. *G* does not take into account regions with zero-counts. We calculated global accessibility on NIH-3T3 fibroblasts (Figure 2A), the proportion of cells in each batch (sh or scr) having global accessibility higher than the median *G* value is enriched in sh fibroblasts (Fisher’s exact test *OR* = 1.616, *p* = 3.751 × 10^*−27*^). Interestingly, values of *G* are inversely correlated to tn5 coverage (Figure 2B). We previously determined that higher coverage is related to replicating cells [1], therefore *G* is higher in G1 cells. This finding is consistent with the fact that deposition of H3K9me3 increases during the S phase, contributing to chromosome condensation [18, 19]. Similarly, enrichment of sh fibrolasts in higher values of *G* is consistent with the fact these cells spend more time in G1 and have a defective S phase. In all, we conclude that the probability of the difference of tn5 and tnH signals in scGET-seq data is a proper proxy to chromatin accessibility.

**Figure 2:**
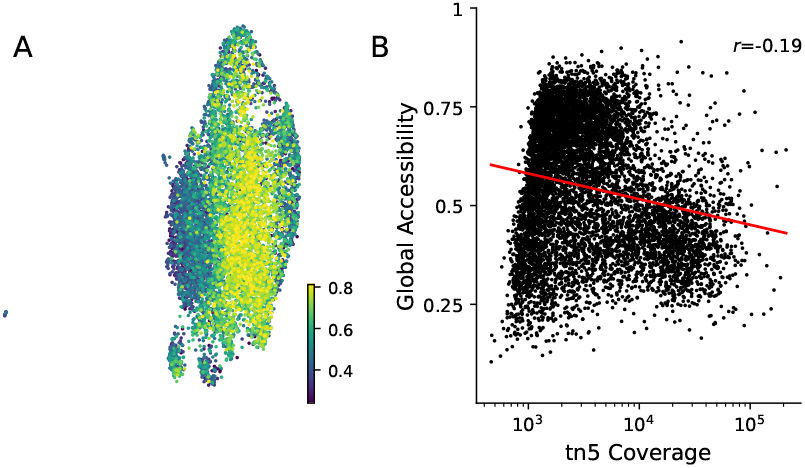
**(A)** UMAP embedding showing cells colored by global accessibility *G* values (Equation 2). **(B)** Scatterplot showing the relation between tn5 coverage and global accessibility *G*. The red line shows the fit of the linear regression between the two variables, together with the Pearson’s correlation coefficient.

### 2.3 Analysis of cell trajectories

Next we wanted to use our definition of accessibility to infer cell states and trajectories. To do so, we exploited the `cellrank` framework [20], in particular its velocity kernel. The main idea of `cellrank`’s velocity kernel is that it uses the values of velocity of each cell and the values of gene expression of neighbour cells to estimate the transition probabilities along the neighbourhood graph. In this kernel, any two measures can be used, as long as one could be thought as the outcome of the other. Therefore, we simply calculate the transition probability posing that the measured net accessibility *A* (defined in Equation 1) is predictive of tn5 counts in neighbour cells. The resulting embedding shows a principal trajectory flowing according to the increase of accessibility (Figure 3A), and analysis of macrostates identifies two major groups (Figure 3B). `cellrank` allows to calculate the correlation of any feature with the probability of being assigned to a macrostate. We used this capability to evaluate the behaviour of pericentromeric regions which are known to be more accessible when Kdm5c is silenced [1, 16]. Correlation evaluated on net accessibility *A* is positive and robustly associated with the inferred trajectory. On the other hand, correlation evaluated on tn5 counts is less robust and, most important, it’s negative (Table 2).

**Table 2:** Correlation of pericentromeric regions used as control to assess chromatin compaction. Values of Accessibility or tn5 signal are correlated to the probability of being part of macrostate 0.

|  | Accessibility |  |  | tn5 |  |  |
| --- | --- | --- | --- | --- | --- | --- |
| | $r$ | pval | qval | $r$ | pval | qval |
| chr2:98660000-98665000 | 0.125 | 3.776e-30 | 3.820e-26 | -0.132 | 4.812e-33 | 7.849e-30 |
| chr2:98665000-98670000 | 0.156 | 8.153e-46 | 4.123e-41 | -0.188 | 3.051e-66 | 9.077e-63 |
| chr4:64495000-64500000 | 0.010 | 3.849e-01 | 8.767e-01 | -0.008 | 4.964e-01 | 8.058e-01 |
| chr6:103645000-103650000 | 0.049 | 9.688e-06 | 4.336e-03 | -0.115 | 1.010e-25 | 7.981e-23 |
| chr9:24540000-24545000 | 0.023 | 3.874e-02 | 4.451e-01 | -0.019 | 8.054e-02 | 2.660e-01 |

**Figure 3:**
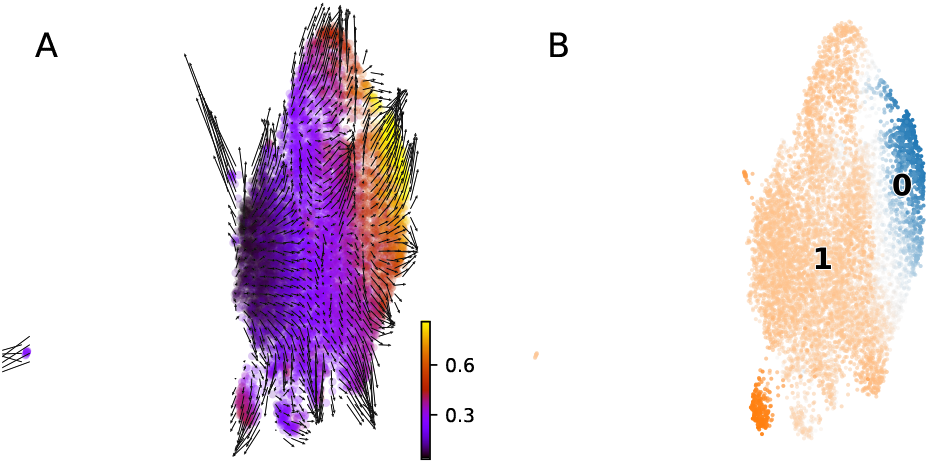
Cell trajectories resulting from analysis of Accessibility. **(A)** Vector field representation of cell trajectories in NIH-3T3 fibroblasts, colored by pseudotime estimated from the velocity kernel. **(B)** UMAP embedding of the same cells colored by cell macrostates, as defined by cellrank.

Negative correlation implies that tn5 decreases while transitioning to macrostate 0, which is in apparent contrast with what previously stated. The average tn5 signal is correlated to coverage (Figure 4A), a known bias in scATAC-seq experiments [21, 8]. Higher coverage in replicating cells can be explained to higher content of DNA [22]. A major consequence of changes in DNA content, and consequently in coverage, is the sparsity of the data, which is higher for resting cells (Figure 4B). Since high number of zeros affects the estimation of averages, we recalculated average tn5 signal excluding zero values, resulting in higher values in resting cells, on the right of the embedding (Figure 4C).

**Figure 4:**
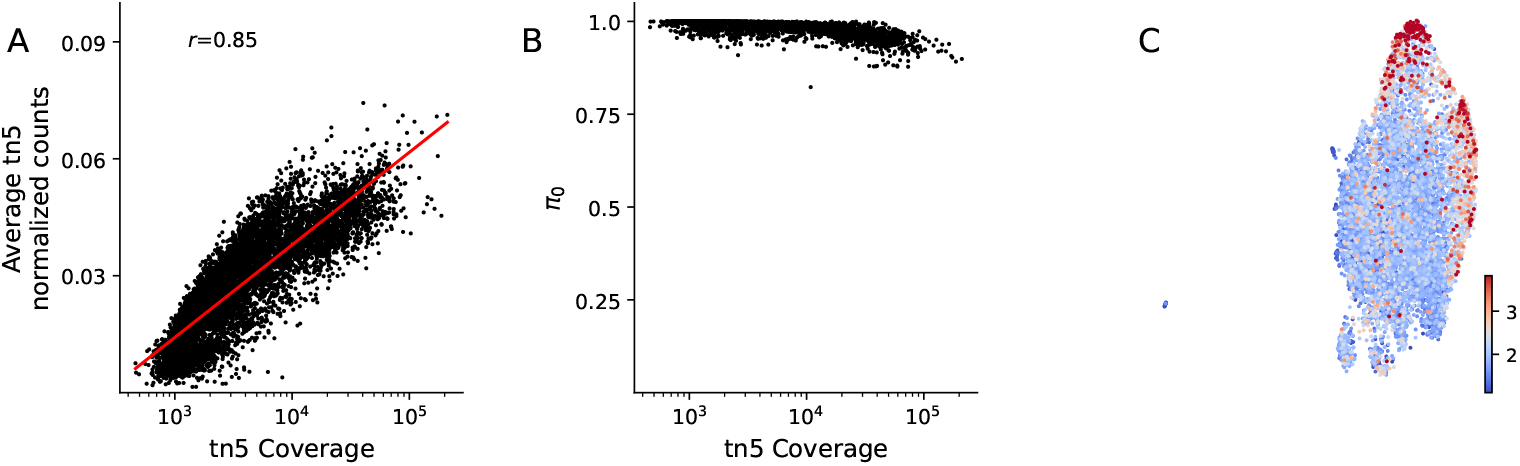
Correlation between coverage and tn5 signal. **(A)** The scatterplot shows the positive correlation between the total per cell coverage and the average amount of tn5 signal retrieved. **(B)** The scatterplot shows the dependency between coverage and data sparsity, measured as the estimated fraction of zero-counts bin (*π*_0_) in ZIP model for tn5 signal. **(C)** UMAP embedding shows cells colored by average tn5 signal excluding zero-counts bins. The signal is higher on the left of the embedding, corresponding to non-cycling cells.

A measure of global accessibility by tn5 alone can be then used to define a `cellrank` kernel based on pseudotime, analogous to CytoTrace scheme [7]. We rescaled tn5 global signal (*tn5 score*) between 0 and 1 and then calculated a tn5 pseudotime as 1*− tn*5 *score*. Similarly, we used global accessibility *G* to define an accessibility pseudotime, then we fitted two Pseudotime kernels to infer cell trajectories and compared them.

Although one should be careful in judging results from the embeddings [23], it is evident that the definition of cell trajectories is noisier when pseudotime kernels are used (Figure 5A,C), compared to the accessibility scheme (Figure 3A). Also, the correlation of tn5 signal in pericentromeric heterochromatin with the probability of being assigned to the appropriate macrostate (2 or 1) is once again negative (Table 3), similarly to what has been highlighted above (Table 2) due to the dependency of tn5 signal on coverage.

**Table 3:**
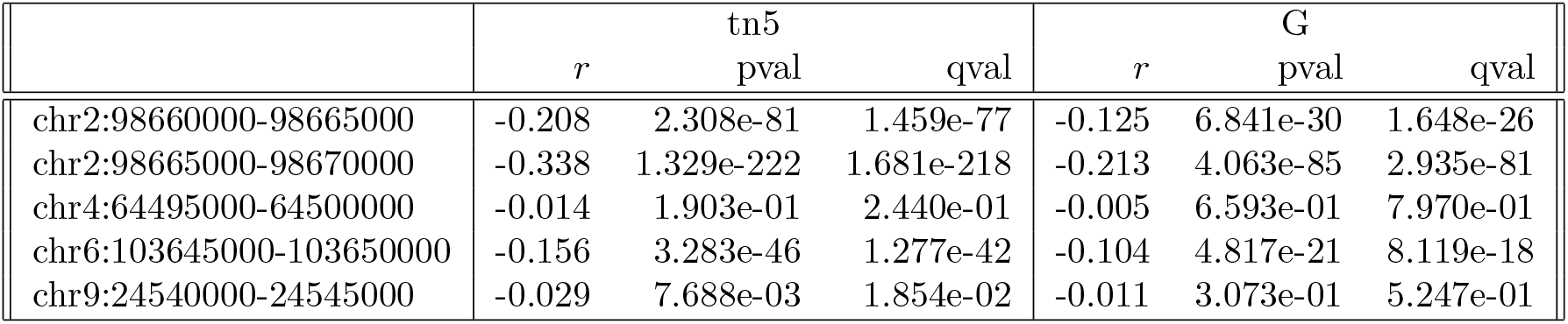
Correlation of pericentromeric regions used as control to assess chromatin compaction. Values of tn5 signal are correlated to the probability of being assigned to macrostates 2 or 1 when a pseudotime kernel is evaluated using tn5 mean values or global accessibility values *G*.

|  | tn5 |  |  | G |  |  |
| --- | --- | --- | --- | --- | --- | --- |
| | $r$ | pval | qval | $r$ | pval | qval |
| chr2:98660000-98665000 | -0.208 | 2.308e-81 | 1.459e-77 | -0.125 | 6.841e-30 | 1.648e-26 |
| chr2:98665000-98670000 | -0.338 | 1.329e-222 | 1.681e-218 | -0.213 | 4.063e-85 | 2.935e-81 |
| chr4:64495000-64500000 | -0.014 | 1.903e-01 | 2.440e-01 | -0.005 | 6.593e-01 | 7.970e-01 |
| chr6:103645000-103650000 | -0.156 | 3.283e-46 | 1.277e-42 | -0.104 | 4.817e-21 | 8.119e-18 |
| chr9:24540000-24545000 | -0.029 | 7.688e-03 | 1.854e-02 | -0.011 | 3.073e-01 | 5.247e-01 |

**Figure 5:**
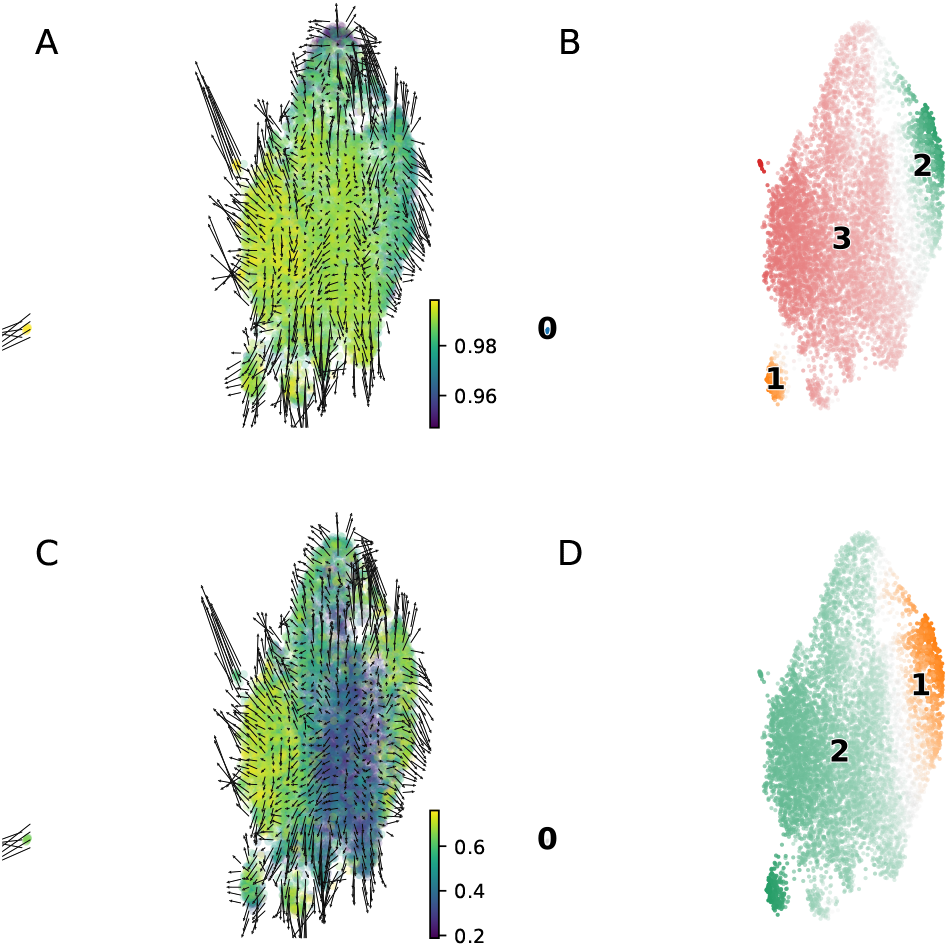
Cell trajectories and macrostates defined using Pseudotime kernels. **(A)** UMAP embedding showing the vector field representation when a pseudotime kernel, defined on tn5 mean accessibility, is used. Cells are colored according to *tn*5_*pseudotime*_. **(B)** Macrostates derived from pseudotime kernel depicted in **A. (C,D)** UMAP embeddings showing results when a pseudotime kernel defined on global accessibility is used.

In all, we found that representing chromatin accessibilty as a net result of tn5 and tnH signals is a simpler solution to handle chromatin modifications in quantitative way and improves the inference on dynamics of epigenome regulation; this measure is not dependent on variations in sequencing coverage due to intrinsic cell properties (*e.g.* cell cycle stauts) and it is more robust than a measure based on accessibility values derived from average signals.

### 2.4 Analysis of iPSC differentiation

In the original publication we analyzed a system in which human induced Pluripotent Stem Cells (iPSC) were derived from skin fibroblasts (FIB) of two unrelated healthy donors, and then differentiated to Neural Precursor Cells (NPC). That system was used to display the controversial framework of chromatin velocity, a term inappropriately introduced for semantic consistency with RNA velocity. We reanalyzed the dataset to compare those results with current framework. We observed a substantial agreement between previous cell groupings and current cell fates (Figure 6), with fibroblasts identified by state 4, iPSC by state 2 and NPC by states 0 and 1.

**Figure 6:**
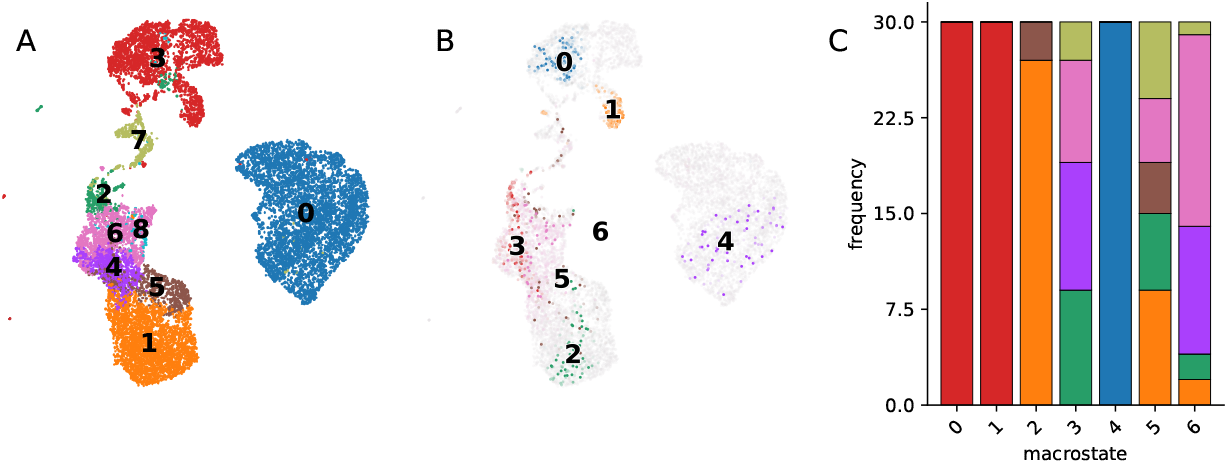
Comparison of previous and current analysis of iPSC model. **(A)** UMAP embedding showing the cell groups identified in [1]. For sake of readibility, the original UMAP embedding has been displayed. **(B)** UMAP embedding projecting currently identified macrostates colored by the absorption probability. **(C)** Histogram showing the proportions of original cell groups in current macrostates.

Willing to understand the link between the two approaches, we analyzed the relation between the likelihood of being a driver region in the velocity model and the region/state correlation as result of the current model (Figure 7A). We observed a positive correlation trend (Pearson’s *r* = 0.299, *p* = 0.0), indicating that regions that were likely scored as putative drivers also have the highest correlation to a macrostate. This suggests that the velocity framework acted as a supercharged linear fitting approach although with limited power, as only a minor fraction of regions could be analyzed. Similarly, we found cell trajectories to be consistent between the two models (Figure 7B, C, Pearson’s *r* = 0.635, *p* = 0.0), suggesting that the previous model, while inappropriate by design, could hint at relevant biology.

**Figure 7:**
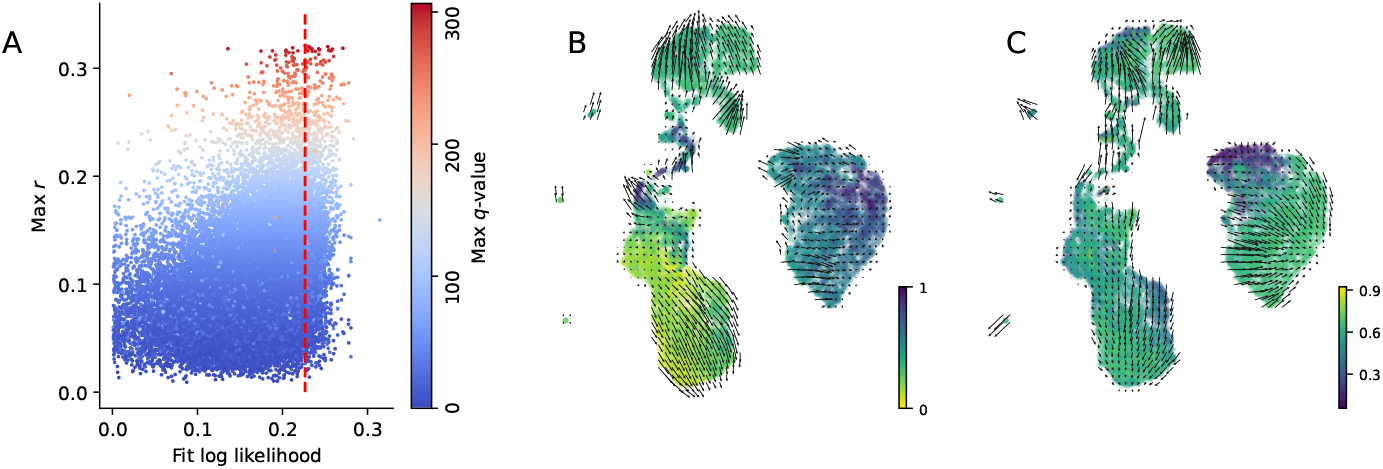
Comparison of previous and current analysis of iPSC model. **(A)** Scatterplot representing the relation between the previously characterized likelihood of velocity model fit on chromatin data and the maximal correlation *r* given by current model to any state. Dots are colored by the minimal *q*-value of the current model. The vertical dashed line indicates the likelihood threshold used in [1]. **(B)** UMAP embedding from the original publication with vector field representation of cell trajectories. Cells are colored by velocity pseudotime. **(C)** UMAP embedding with vector field representation of cell trajectories returned by current analysis. Cells are colored by global accessibility score.

In previous analysis, we identified 1,703 regions that were considered drivers (velocity drivers, VD), such regions were also associated to specific cell groups. We compared the correspondence of VD to cellrank drivers (CD), looking for positive associations with *q <* 0.01. We found 918 VD/CD pairs for which the cell group/macrostate correspond. In addition 317 VD were not associated to any cell group, and 414 VD are not associated to the correct macrostate (*F*_1_ = 0.700).

## 3 Discussion

Single cell GET-seq was introduced to study genome and epigenome properties from the same cells at high throughput. It relies on the combined usage of two transposases: tn5 and tnH, the latter with a higher affinity for heterochromatin. The original analysis workflow is based on the evidence that analysis of scATAC-seq data on predefined regions is comparable with *de novo* discovery of peak locations [24]. Nevertheless, read counts from compact chromatin should also be acquired to analyze scGET-seq data; for this reason, the original workflow relied on the acquisition of four data matrices, one for DHS rich and one for DHS poor regions for each transposase, and their projection in a common latent space by matrix tri-factorization [2]. While effective, that approach requires a prior knowledge of regulatory regions for the organism under study, which is not necessarily true for non-human or non-model organisms. An agnostic approach, based on genome binning, is more generalizable and can still be integrated with prior knowledge at post-processing time. In this paper we propose a strategy based on fixed-size windows, analogous to other analysis frameworks [11], which has two major advantages: first, it allows to represent data using a single tensor which we show can be efficiently factorized to obtain a low-dimensional embedding of the single cells; second, read counts in each bin do not depend on the bin size and can be easily used to built a statistical representation of the data at the single cell level.

Among the possible approaches for tensor factorization, we chose Tensor Train Decomposition, and we showed that it results in expressive data representation, conserving biological information. To model read counts from two transposases within the defined bins, we introduced a new approach that defines accessibility as the likelihood of observing a specific difference in read counts between the two transposases. We show that such measure can be applied to derive a global score of genome accessibility, which can be used to characterize cell phenotypes. Global accessibility is not dependent on total coverage, contrary to a central measure based only on tn5 signal identified in scATAC-seq experiments.This approach to scGET-seq data overcomes the limitations of chromatin velocity [1] and allows more robust identification of cell states and trajectories. Velocity frameworks are not suited to study chromatin dynamics, in that they make assumptions that do not hold for scGET-seq data. However, we show that such framework did result in credible results, possibly due to their capabilities to fit given data. Accessibility, as defined in this paper, is a definition that makes very few assumptions (*e.g.* the independence of tn5 and tnH activity) and applies data transformation in a way that is consistent across cells. Nonetheless, we anticipate that the field of study would greatly benefit from the description of chromatin dynamics by a mechanistic model that takes into account the rate of compaction and/or relaxation.

Overall, in this paper we devised a novel statistical strategy that is able to robustly identify closed and open chromatin at the single cell level, and showed that this metric can be used to describe global cell states, such as the chromatin alterations subjected by Kdm5c, known for its role in altering global chromatin compaction [15]. Moreover, we showed how to reduce the dimensionality and embed the combined information from tn5 and tnH, using the simple and reliable method of tensor decomposition. We didn’t introduce alternative strategies to treat genomic data, such as mutations and/or copy number alterations, considering the increasing number of strategies could easily be applied to scGET-seq [25, 26]. Concluding, we foresee that methodologies described in this paper will be widely used to analyze states of chromating compaction, as well as their regulation and dynamics.

## 4 Methods

Unless stated differently, all the analysis were performed using scanpy v1.8.2 [27], scvelo v0.2.4 [5], cellrank v1.5.1 [20], schist v0.7.14, [12] and tensorly v0.7.0 [28].

### 4.1 Fit of count data

Poisson’s distribution is characterized by a single parameter *λ* that is simply estimated as

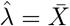

where *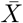* is the average of bin counts for a cell, excluding mitochondrial reads. Zero Inflated Poisson distribution is defined by two parameters *λ*, the expected counts in a bin, and *π*_0_, the expected number of empty bins. We fitted ZIP we using Maximum Likelihood Estimator (MLE) to perform likelihood ratio test and verify that ZIP is indeed more suited to describe our count data. Given the average cell per-bin read counts (excluding mitochondrial reads) *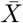* and the cell average number of empty bins*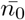*, the square root of sample variance 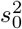 is

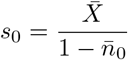

and the MLE for *λ* is

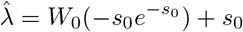

where *W*_0_ is the principal solution of Lambert’s W function (evaluated using `scipy.special.lambertw`). Then the MLE for *π*_0_ is

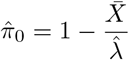

For practical purposes (speed and numerical stability), instead, we used the Method of Moment Estimator (MME), so that the expected counts is defined as

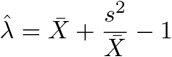

where *s*^2^ is the sample variance, and the fraction of zeros is

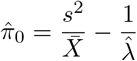

To avoid negative values of *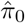* that happen when *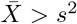* we truncated *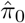* [17], hence

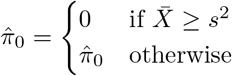

and

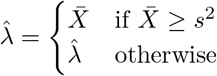

Log-likelihood ratio test was performed on each cell using the fitted distributions, the *p*-value was calculated using *χ*^2^ distribution with 1 degree of freedom.

### 4.2 Analysis of Caki-1/HeLa cell mixture

Raw reads were downloaded from ArrayExpress (ID: E-MTAB-9650) and processed using default scGET-seq pipeline [29]. Read counts were produced using 5kb bins over hg38 reference genome. Original processed data used in the original paper [1] were used to annotate cell identities and for matrix tri-factorization embedding. After fitting ZIP distribution, we removed cells having a total coverage (as sum of tn5 and tnH coverage) lower than 2,000. We identified low-quality regions as those having zero counts in both tn5 and tnH in less than 90% of the cells.

Count values for tn5 and tnH were normalized per total coverage (*i.e.* the sum of tn5 and tnH coverages) and transformed by `log1p`. The normalized counts were used to build the tensor subsequently decomposed by TTD using a rank [1,200,1]-decomposition, so that we obtained three core matrices *W, E, F*. Matrix *W* has shape (2, 1) and can be used to weight the relevance of each enzyme counts to the decomposition. Matrix *E* has shape (*n_cell_,* 200) and it is considered the low-dimensional embedding of cells. Matrix *F* has shape (200, *n_features_*) and it is currently discarded. Normalized tn5 counts were also used to calculate PCA embedding.

For each embedding, we derived the *k* NN graph using 20 components and setting the number of neighbors equal to*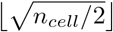*. We identified hierarchical cell groups by nested Stochastic Block Models implemented in `schist`, then we chose the highest level in hierarchy that produces a split in at least two groups. Clustering performance on each embedding was evaluated using `scikit-learn` v0.24.2 [30].

### 4.3 Analysis mouse fibroblasts

Raw reads were downloaded from ArrayExpress (ID: E-MTAB-9651) and processed using default scGET-seq pipeline [29]. Read counts were produced using 5kb bins over mm10 reference genome. After fitting ZIP distribution, we removed cells having a total coverage (as sum of tn5 and tnH coverage) lower than 2,000. We identified low-quality regions as those having zero counts in both tn5 and tnH in less than 90% of the cells.

Count values for tn5 and tnH were normalized per total coverage (*i.e.* the sum of tn5 and tnH coverages) and transformed by `log1p`. The normalized counts were used to build the tensor subsequently decomposed by TTD using a rank [1,200,1]-decomposition.

Once ZIP laws were fitted for each enzyme, we calculated local accessibility *A* for each bin according to Equation 1. Next we calculated global accessibility *G* (Equation 2). To proceed with `cellrank` analysis, we smoothed the normalized tn5 counts and the matrix *A* using the *k* NN graph. The cr.tl.kernels.VelocityKernel was initialized using default parameters and transition probabilities were calculated using cosine distance. The kernel was then regularized using a connectivity kernel (0.8*v* + 0.2*c*). Cell macrostates were identified using GPCCA estimator (`cr.tl.estimators.GPCCA`) using default parameters. Analysis of correlation between pericentromeric regions and macrostate assignment was performed using the `return drivers` function, using values of *A* or normalized tn5 counts.

`cr.tl.kernels.PseudotimeKernel` class was used to analyze cell trajectories using average tn5 counts or global accessibility. Pseudotime kernels were initialized using default parameters, followed by GPCCA estimation of cell macrostates. We choose to extract the lowest number of states that did not raise numerical errors due to matrix irreducibility.

### 4.4 Analysis iPSC

Raw reads were downloaded from ArrayExpress (ID: E-MTAB-10218) and processed using default scGET-seq pipeline [29]. Read counts were produced using 5kb bins over hg38 reference genome. After fitting ZIP distribution, we removed cells that were not analyzed in [1], retaining 12,800 single cells. We identified low-quality regions as those having zero counts in both tn5 and tnH in less than 90% of the cells.

Accessibility was calculated as described in the previous section and similarly it used to initialize a `cellrank` Velocity kernel. UMAP embeddings were retrieved from original publication, as well as the fit likelihoods for the velocity model.

## Acknowledgements

We thank Giulio Caravagna, Riccardo Bergamin and Salvatore Milite at University of Trieste, Paolo Provero at University of Turin and all the members of the COSR laboratory for discussions and/or critical reading of the manuscript. This work was partially supported by the AIRC/CRUK Accelerator Award “Single Cell Cancer Evolution in the Clinic” A26815 and by the AIRC/CRUK/AECC Accelerator Award “Early Cancer Research Initiative Network on MBL and MGUS models (ECRIN-M3)” A29370.

